# IROP: Ionic Regulation of Permeation for High-Yield AAV Purification

**DOI:** 10.64898/2026.01.16.699902

**Authors:** Deepraj Sarmah, Scott M. Husson

## Abstract

Traditional filtration methods for separating host cell proteins and DNA from adeno-associated viral capsids often result in substantial yield losses. Here, we introduce a new high-yield purification strategy termed Ionic Regulation of Permeation (IROP) for removing major impurities within a single unit operation. Rather than relying on static size exclusion or binding interactions, IROP dynamically modulates AAV transport within a nanoporous membrane. At low ionic strength, AAV permeation was negligible, while high ionic strength markedly increased the sieving coefficient, revealing an electrostatic gating mechanism. AAVs retained under low-salt conditions could not be efficiently recovered by a static high-salt wash, indicating steric-electrostatic trapping within membrane pores. In contrast, high-salt filtration enabled high recovery. Leveraging this mechanism, a two-step IROP process, involving low-salt impurity clearance followed by high-salt AAV recovery, achieved greater than 90% AAV recovery with a greater than 95% reduction in HCPs and host cell DNA.

## Introduction

Adeno-associated virus (AAV) has become a leading platform for *in vivo* gene therapy, with multiple approved products and hundreds of ongoing clinical trials^1,2^. Yet, the broad deployment of AAV-based therapies remains constrained by the high cost and complexity of manufacturing, with downstream purification representing a significant bottleneck^3^. Purification must efficiently separate full viral capsids from a complex milieu of product-related impurities (e.g., empty capsids) and process-related impurities, particularly host cell proteins (HCPs) and host cell DNA. Residual HCPs can elicit immune responses, and residual DNA presents potential oncogenic risk, making their stringent removal essential for product safety and regulatory acceptance^4^.

Industry-standard downstream processes commonly incorporate tangential flow filtration (TFF) along with other unit operations. TFF is highly scalable and widely used for buffer exchange and concentration. However, when TFF is used for purification, the operation relies on passive, size-based discrimination and therefore faces an inherent trade-off: membranes sufficiently tight to retain AAVs often promote fouling, aggregation, and poor clearance of similarly sized impurities, such as host DNA fragments, ultimately resulting in variable and substantial product losses of 20-40%^5^. Moreover, there is a trade-off between AAV loss and impurity clearance. Membranes with smaller pores (100 kDa) retain more AAV but remove fewer impurities^6^, while membranes with larger pores (500 kDa) remove more impurities alongside a substantial loss of AAV^7^.

This limitation reflects an oversimplified view of membranes as purely steric sieves. For nanoporous membranes, there is also the potential to exploit the electrostatic effects resulting from different ionic strengths and pH, which can often dominate steric effects. The concept of electrostatic gating in charged nanopores demonstrates that the functional pore size can be modulated by ionic strength^8^. At low ionic strength, an electrostatic double layer extends into the pore, generating a repulsive energy barrier that excludes co-ions, effectively “closing” the pore. At high ionic strength, charge screening collapses the double layer, “opening” the pore for sterically permitted transport^9,10^. Although well-established in membrane science, this gating behavior has not previously been harnessed to create a tunable, selective separation method for a complex biologic like AAV. Membranes, such as polyethersulfone (PES), have been shown to exhibit a high negative zeta potential at low ionic concentrations, which can be significantly neutralized by increasing the ionic concentration or lowering the pH^11,12^. At low ionic strength, strong electrostatic gating occurs: the highly negative membrane surface charge repels AAV particles at pH values above their isoelectric point (>6.3)^13^, resulting in high AAV rejection. At higher ionic strengths, charge screening reduces the membrane zeta potential and weakens electrostatic gating, resulting in lower AAV rejection.

In previous work, we have exploited steric interactions of AAV with membrane pores to demonstrate the separation of full and empty capsids^14^. Here, we leverage both steric and electrostatic properties in a distinct bioprocessing strategy we term **I**onic **R**egulation **o**f **P**ermeation (IROP). We demonstrate that a standard PES membrane can be dynamically switched between an AAV-retentive state at low ionic strength and an AAV-permeable state at high ionic strength. A key mechanistic insight is that AAVs retained under low-salt conditions become sterically and electrostatically trapped within the pore network, a behavior that enables highly selective separation. Unlike chromatographic or binding-based methods, IROP operates purely through dynamic modulation of transport within a convective membrane system. This capability allows IROP to decouple yield from purity, achieving greater than 90% AAV recovery while simultaneously reducing HCPs and host cell DNA by more than 95%. This work establishes a new purification paradigm based on tunable, charge-regulated permeation, offering a fundamentally different mode of separation for viral vectors and other nanoscale biologics.

## Results

### AAV permeation through 30 nm membranes is strongly regulated by ionic strength

To evaluate how ionic conditions affect AAV transport through a 30 nm PES membrane, we measured sieving coefficients by filtering half the volume of a purified AAV solution in three buffers: a low-salt buffer (composition in Methods), a buffer containing 100 mM MgCl_2_, and a buffer containing 250 mM NaCl. Filtration experiments were conducted at nominal pressures of 35, 70, 105, and 140 kPa.

Under low-salt conditions, AAV transport was minimal across all pressures. The sieving coefficients were generally <0.01, indicating near-complete rejection (Fig. 1a, 1d). Values increased slightly with increasing pressure, from 7.25×10^-3^ at 35 kPa to 1.25×10^-2^ at 140 kPa, suggesting that an electrostatic energy barrier impedes AAV passage through the pores at low ionic strength, and increased convection partially overcomes this barrier.

**Figure 1:**
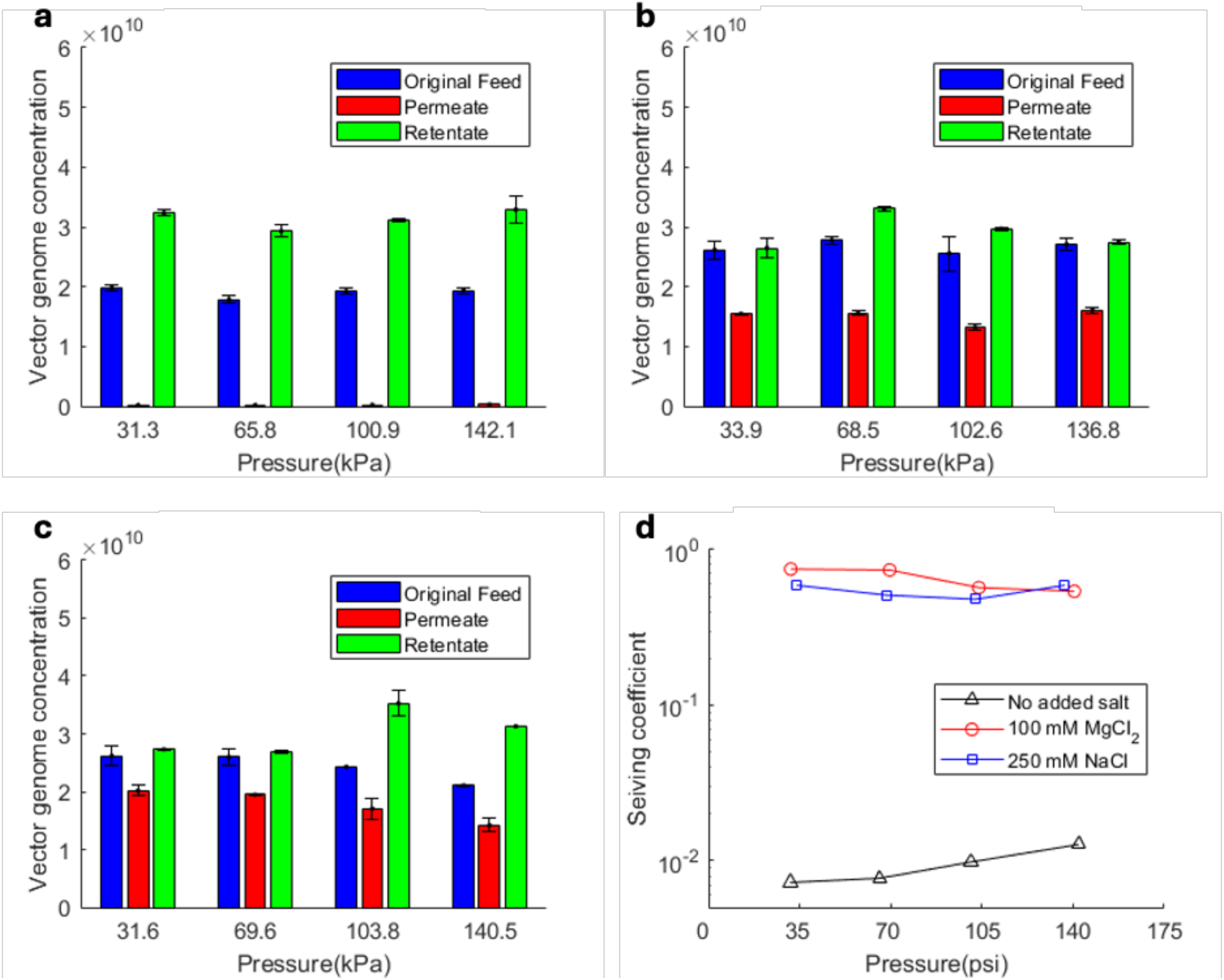
Measured AAV2 vector genome concentrations in the original feed, permeate and retentate after diafiltration with (a) low salt buffer, (b) buffer containing 250 mM NaCl, and (c) buffer containing 100 mM MgCl_2_. (d) Pressure-dependent AAV2 sieving coefficients under the same buffer conditions.

Elevating the ionic strength led to a marked increase in AAV permeation. At 250 mM NaCl and 100 mM MgCl_2_, the sieving coefficients rose to ∼0.6 (Fig. 1b, 1d) and ∼0.7 (Fig. 1c, 1d), respectively, and were insensitive to pressure. These observations are consistent with electrostatic gating: low ionic strength “closes” the pores by extending electrostatic double layers, whereas high ionic strength screens these interactions and “opens” the pores, allowing AAV transport. The disappearance of pressure dependence at high ionic strength further supports ionic gating as the dominant mechanism.

We performed the same measurements using hemoglobin (0.5 mg/mL) as a model HCP and using diluted cell lysate (as described in Methods), filtered at 70 kPa. In contrast to AAV, hemoglobin and HCPs permeated readily under low ionic strength conditions (Fig. 2a-c). These contrasting behaviors, characterized by high AAV rejection and high impurity permeation, suggest an opportunity for purification based solely on ionic strength modulation.

**Figure 2:**
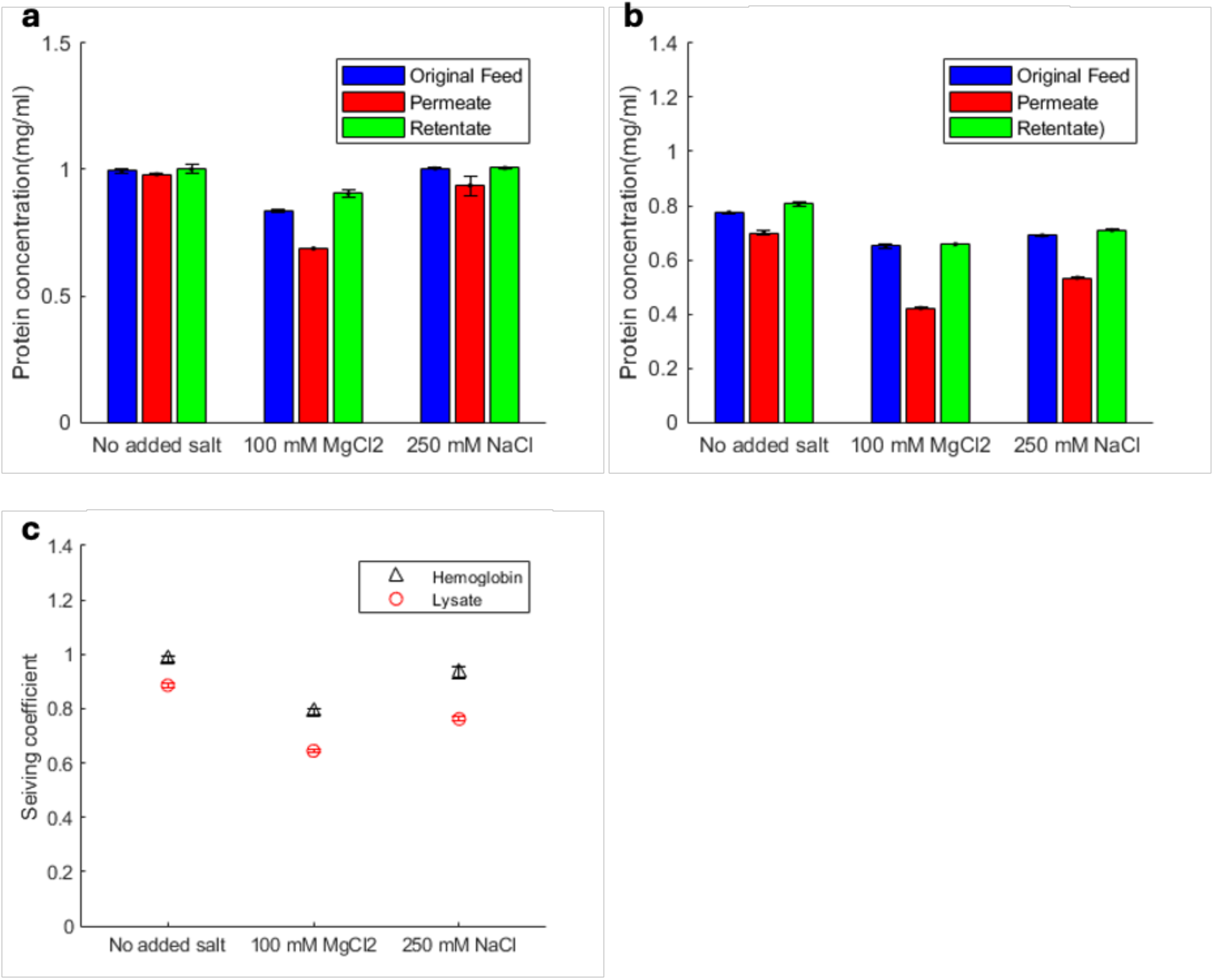
(a) Measured protein concentration for (i) hemoglobin and (ii) lysate HCPs in the original feed, permeate and retentate after diafiltration at 70 kPa in low salt buffer, and buffers containing either 250 mM NaCl or 100 mM MgCl_2_. (b) Measured sieving coefficients of hemoglobin and lysate HCPs under the same buffer conditions.

### Recovery experiments reveal steric-electrostatic intra-pore trapping of AAV

To leverage this behavior and elucidate the retention mechanism, we designed a two-step process for HCP and DNA removal, followed by the subsequent recovery of AAV.

1. **Impurity clearance (low ionic strength filtration):** Two-stage diafiltration was performed at a low pressure (∼15-30 kPa) in a low-salt buffer. As expected, the permeate fractions contained only 2-3% of the initial AAV, as measured by qPCR (Fig. 3a, b). Surprisingly, the retentate contained only 8-12% of the initial AAV. Because the permeate carried very little AAV, the low recovery in the retentate indicates that AAV is not simply accumulating on the membrane surface. It must be entering the membrane structure and becoming retained within the pore network. The first diafiltration stage removed 75-80% of HCP and 80-85% of DNA. The second stage removed an additional 8-10% of these impurities.
2. **AAV recovery (high ionic strength filtration):** After diafiltration, we tested two recovery methods under high-salt conditions: (i) washing the retentate side with 250 mM NaCl without filtration, and (ii) adding 100 mM MgCl_2_ to the retentate and filtering it through the membrane.

**Figure 3:**
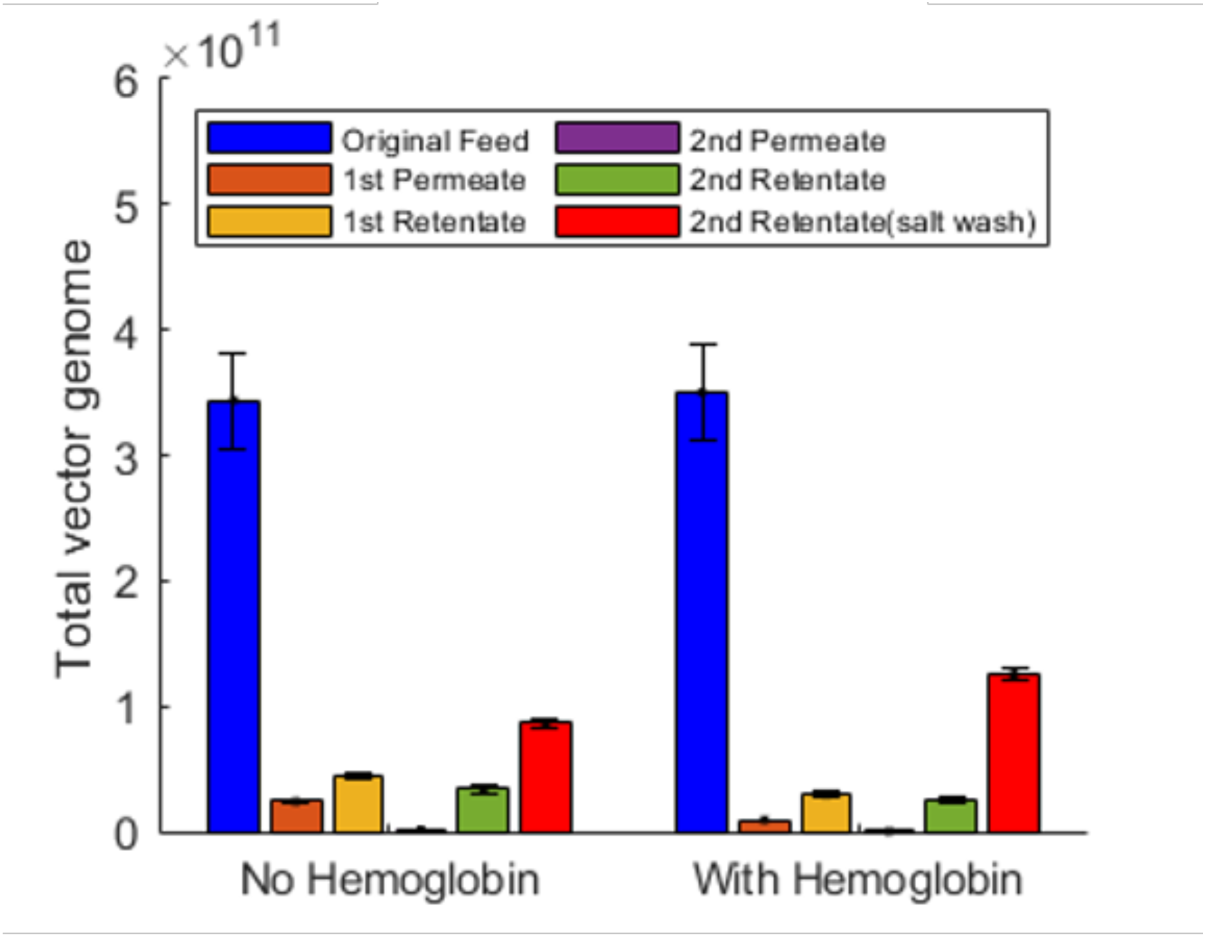
Measured AAV2 total vector genomes for the original feed; first permeate and retentate fractions collected in low salt buffer; second permeate and retentate fractions collected in low salt buffer; and the final retentate (salt wash) collected with high salt buffer (250 mM NaCl).

Washing the retentate side recovered only ∼25-35% of the AAV, even with vigorous vortexing (Fig. 3, 2^nd^ retentate salt wash). This low recovery yield rules out simple surface deposition as the dominant mechanism. In contrast, filtration with the high-salt buffer recovered ∼75% of the AAV (Fig. 4a). This disparity between washing and filtration is consistent with steric-electrostatic intra-pore trapping: AAVs appear to be convectively driven into confined pore regions at low ionic strength, requiring both charge screening and convective transport to achieve release.

**Figure 4:**
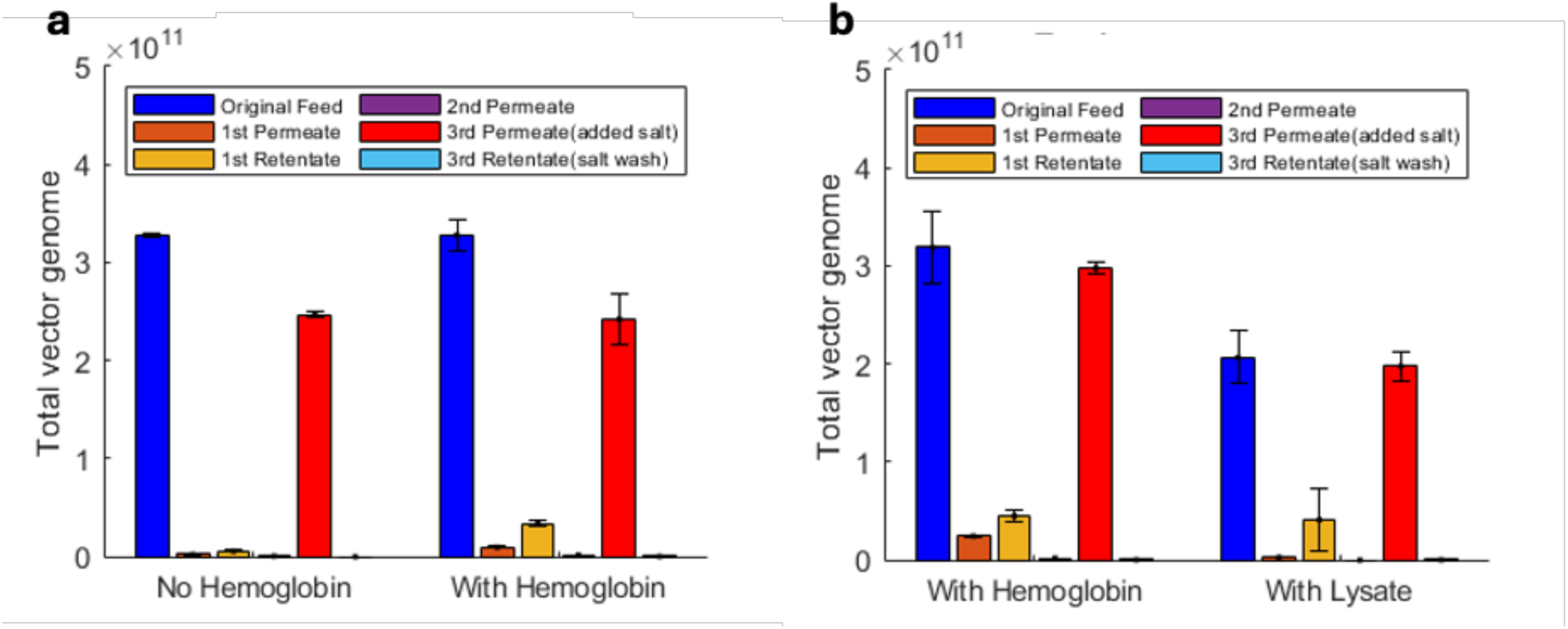
Measured AAV2 total vector genomes in the original feed, first permeate and retentate fractions collected in low salt buffer; second permeate collected in low salt buffer; permeate collected after addition of high salt buffer; and the final retentate wash collected with salt solution. Results are shown for (a) 100 mM MgCl_2_, using an AAV2 solution without added hemoglobin and with added hemoglobin and (b) 150 mM MgCl_2_, using an AAV2 solution with added hemoglobin and with added lysate (diluted 1:20).

We refer to this purification strategy as **I**onic **R**egulation **o**f **P**ermeation (IROP). In IROP, AAV is confined within the membrane pores during low-salt filtration in the first step and released during high-salt filtration in the second step. Further investigation of IROP using a higher concentration of MgCl_2_ (150 mM) improved AAV recovery yield to 93-95% (Fig. 4b). Under these conditions, the remaining hemoglobin/HCPs and DNA in the final product were 2-7% and 5-7%, respectively. The experiment with lysate was performed twice to confirm repeatability.

### Scanning electron microscopy reveals large, unfouled surface pores

Scanning electron microscopy of the membrane surface revealed pore sizes of approximately 200-500 nm (Fig. S1), substantially larger than the nominal 30 nm pore rating specified by the manufacturer and larger than AAV capsid diameters (∼25-30 nm). Imaging of low and high ionic strength filtrations showed no detectable accumulation of AAV particles on the membrane surface, indicating that the surface remained unfouled under all conditions examined. SEM imaging of both sides of the membrane showed similarly large and open pores.

## Discussion

What mechanism underlies the ability of the membrane to retain AAV while simultaneously allowing HCPs and DNA to pass? The behavior most likely arises from the differential interactions of these species within the nanopores. In contrast to AAVs, HCPs are generally significantly smaller, less spherical, and more flexible than AAVs. These characteristics allow them to navigate the pore’s electrostatic landscape more freely. Host cell DNA, as a large anionic polymer, adopts an extended conformation within the pore walls^15,16^, and the pressure-induced convective flow during low-salt diafiltration is sufficient to sweep both impurity classes through the membrane. At the same time, the IROP mechanism ensures that AAVs remain trapped.

The mechanism of AAV capture within the membrane at low ionic strength can be rationalized by considering the interplay among particle size, pore geometry, and electrostatic interactions. Importantly, this process is mechanistically distinct from chromatographic bind-and-elute behavior. At the operating pH, both the PES membrane and the AAV capsid carry net negative charge; thus, attractive binding is absent, and the membrane cannot retain AAV through an affinity-like mechanism. Instead, retention arises from repulsive electrostatic interactions that become increasingly restrictive within confined pore regions. AAV capsids are rigid, ∼25 nm diameter, near-spherical particles with complex surface charge distributions that make them highly sensitive to electrostatic double-layer forces in confined environments. Although the membrane surface appears sterically permissive, with pores much larger than the AAV diameter, the strong ionic-strength dependence of AAV sieving suggests that retention occurs within more confined regions of the pore network where electrostatic repulsion is higher. As ionic strength increases, charge screening reduces these interactions, allowing AAV to pass through regions of the membrane that were inaccessible under low-salt conditions.

The SEM observations refine this mechanism. Imaging revealed large surface pores on both sides of the membrane, with no visible accumulation of AAV at the membrane surface. These findings rule out alternative explanations such as surface fouling or hydrodynamic pore-mouth blockage, in which particles accumulate at pore entrances due to converging streamlines rather than entering the pores. The absence of AAV at the surface is inconsistent with a purely surface-based retention mechanism and instead indicates that AAV particles enter the pore network before becoming trapped.

Because both membrane surfaces have similarly large pores, a standard asymmetric pore structure model is insufficient. One possibility is that the membrane features an internal hourglass-shaped architecture, characterized by large pores on each surface and a constricted zone located deeper within. Under low ionic strength, electrostatic gating at these narrowed regions could restrict AAV passage. Hourglass-type^17–19^ or bidirectionally tapered nanopore^20^ architectures have been reported in other filtration materials, as revealed through cross-sectional or tomographic imaging, which would be a valuable direction for future work.

Other lines of evidence further support this mechanistic interpretation. First, hydrodynamic blockage alone cannot account for the strong ionic-strength dependence of AAV sieving. Pore-mouth jamming for these pore sizes is expected to depend more strongly on the presence of larger macromolecules, such as HCPs and DNA, than on salt concentration^21–23^. In contrast, we observe no significant difference between experiments conducted with or without hemoglobin or lysate, whereas a greater than 50-fold increase in AAV permeability is observed when transitioning from low to high ionic strength. Second, hydrodynamic blockage would be expected to impede the passage of smaller species as well, yet hemoglobin, HCPs, and host DNA permeate efficiently under low-salt conditions. Third, the inability of the static high-salt washing to recover AAV argues against surface-level blocking; instead, recovery requires both charge screening and convective transport through the membrane to release particles trapped within confined pore regions. Although AAV recovery is not fully quantitative, the available evidence indicates that the unrecovered fraction does not arise from surface fouling or pore-mouth blockage, as no capsid accumulation is visible on the membrane surface. We have not yet evaluated cycling behavior, and future studies will be needed to determine whether repeated operation introduces any irreversible retention or fouling effects.

Taken together, these results support a model in which steric–electrostatic interactions within the pore interior, not surface fouling or hydrodynamic entrance effects, govern selective AAV trapping during IROP. By exploiting the distinct physicochemical properties of AAV, HCPs, and host cell DNA, IROP achieves AAV recoveries of over 90% with greater than 95% clearance of HCPs and host cell DNA in a single, tunable unit operation. This performance exceeds current industry norms for filtration-based purification, suggesting that IROP may serve as a generalizable platform technology for purifying other viral vectors, virus-like particles, or nanoscale biologics whose sizes and surface charge profiles permit controlled intra-pore interactions.

While the collective data strongly indicate that intra-pore steric and electrostatic effects dominate AAV retention, we acknowledge that minor contributions from pore-mouth obstruction or restricted surface transport cannot be excluded entirely. Future imaging studies, particularly cross-sectional or 3D characterization, will be necessary to resolve the internal pore architecture and the distribution of AAV within the membrane structure. Additionally, the potential effects of impurity size and charge heterogeneity on IROP performance are acknowledged but not evaluated in this work.

## Materials and Methods

### AAV and Cell Lysate Source

Adeno-associated virus serotype 2 (AAV2-CAG-GFP-Stuffer; Catalog #449B459-2-500), containing a recombinant genome encoding eGFP under the control of a CAG promoter, was obtained from Virovek (Houston, TX, USA). Empty AAV capsids (AAV2-Empty; Catalog #449B000-2-500) were procured from the same supplier. The manufacturer-supplied clarified lysate from Virovek was used as the feed material for all filtration experiments after prefiltration through a 1 µm regenerated cellulose membrane (Tisch Scientific, Cleves, OH; SKU 10410012) to remove large cellular debris and organelles.

### Membranes and Filtration Systems

All IROP experiments were performed using polyethersulfone (PES) ultrafiltration membranes with a nominal pore size of 30 nm (Sterlitech, Auburn, WA). Filtrations were conducted in a 50 mL direct-flow stirred ultrafiltration cell (Amicon®, MilliporeSigma, Burlington, MA) equipped with a pressure regulator and a magnetic stirrer operating at 200 rpm to minimize concentration polarization. Experiments were operated at ambient temperature, and a compressed air cylinder supplied the required pressure.

### Sieving Coefficient Analysis

The observed sieving coefficient (S_obs_) for AAV was determined using a 1:1 mixture of full and empty AAV particles. Filtrations were performed at 138 kPa. AAV concentrations were measured in the feed (C_F_), permeate (C_P_), and final retentate (C_R_).

Because the concentration in the retentate increases during filtration, the average AAV concentration in the retentate over the course of the experiment (⟨C⟩_R_) was approximated by the arithmetic mean of the initial feed concentration and the final retentate concentration:

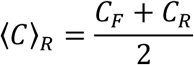

The observed sieving coefficient was then calculated as:

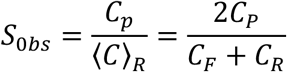

This formulation accounts for the increase in solute concentration in the retentate during filtration and provides a consistent estimator of the permeation behavior under the operating conditions used.

### Feed Preparation for Impurity Clearance Experiments

Purified AAV samples or cell lysate were diluted in a low-salt buffer comprising 20 mM tris-ascorbic acid and 1 mM EDTA at pH 7.8. For experiments using purified AAV, 10 µL each of purified full AAV (AAV2-CAG-GFP-Stuffer) and purified empty AAV (AAV2-empty) were diluted into 20 mL of the low-salt buffer.

In experiments where hemoglobin was added as a model host cell protein, 10 mg of hemoglobin was added to the purified AAV solution in 20 mL of low-salt buffer, resulting in a hemoglobin concentration of 0.5 mg/mL. For experiments using clarified lysate, 1 mL of lysate was diluted into 20 mL of the low-salt buffer.

## Two-Step IROP Purification Procedure

### Low-salt HCP/DNA Clearance (Diafiltration)

A diafiltration cycle consisted of passing approximately 90% of the feed volume (using pre-clarified lysate) through the membrane, followed by replenishment of the feed reservoir with fresh buffer to restore the initial working volume. This procedure was repeated for a total of two diafiltration cycles. Permeate generated during diafiltration, containing mainly host-cell proteins and DNA, was collected.

### High-salt AAV Recovery

Following diafiltration, AAV was recovered using a high-salt buffer. Initial static salt wash experiments utilized a buffer comprising 250 mM NaCl, 20 mM tris-ascorbic acid, and 1 mM EDTA at a pH of 7.8. In these experiments, the retentate side of the membrane was washed with the high-salt buffer, and the solution was collected and analyzed to quantify recovered AAV.

Subsequent IROP experiments used a high-salt buffer containing 100 mM MgCl_2_, 20 mM tris-ascorbic acid, and 1 mM EDTA at a pH of 7.2. In these experiments, the retentate side was washed with the high-salt buffer, and the solution was passed through the membrane under pressure (∼140 kDa). The resulting permeate was collected and analyzed to quantify recovered AAV.

For experiments yielding the highest AAV recovery (>93%; see Table S1), the same procedure was followed using a higher MgCl_2_ concentration (150 mM) in the buffer solution at a pH of 6.8-7.

In IROP optimization experiments (Figs. 3, 4b) were conducted using purified AAV solutions with and without hemoglobin as a model HCPs. After achieving the effective removal of hemoglobin and a high recovery of AAV, the method was applied to purify AAV from a diluted lysate.

Samples from the feed, low-salt permeate, and high-salt permeate were analyzed for AAV genome titer, HCP concentration, and host cell DNA to determine yield and impurity clearance.

## Analytical Methods

### AAV Genome Titer (qPCR)

AAV genomes were quantified by qPCR using primers targeting the inverted terminal repeat (ITR) sequences (forward: 5-GGA ACC CCT AGT GAT GGA GTT-3; reverse: 5-CGG CCT CAG TGA GCG A-3; Integrated DNA Technologies, Coralville, IA, USA). Standard curves were generated using AAV reference material (Addgene, Watertown, MA, USA; catalog number 59462-AAV2).

### Total capsids (ELISA)

Total capsid concentrations were measured using an AAV2-specific ELISA kit (Progen Biotechnik (Heidelberg, Germany; catalog number: PRAAV2R). Absorbance was read on a Synergy H1 microplate reader (Aligent BioTek, Winooski, VT, USA).

### Host-cell Protein Assay

HCP concentrations were quantified using a Bradford protein assay with BSA standards. Absorbance was measured using the Synergy H1 microplate reader.

### Host-cell DNA (PicoGreen)

Host cell dsDNA was quantified using the HyperLight™ PicoGreen dsDNA assay (APExBIO, Houston, TX, USA) and read on the Synergy H1 microplate reader.

### Statistical Analysis

Measurements from all filtration experiments were performed in triplicate, except for lysate filtration experiments, which were conducted in duplicate. Data are presented as mean ± standard deviation.

## Author Contributions

Conceptualization, D.S. and S.M.H.; methodology, D.S. and S.M.H.; validation, D.S. and S.M.H.; formal analysis, D.S. and S.M.H.; investigation, D.S.; resources, S.M.H.; data curation, D.S.; writing—original draft preparation, D.S.; writing—review and editing, S.M.H.; visualization, D.S.; supervision, S.M.H.; project administration, S.M.H.; funding acquisition, S.M.H. All authors have read and agreed to the published version of the manuscript.

## Institutional Review Board Statement

The study was approved by the Institutional Biosafety Committee of Clemson University (IBC2023-0102, effective from February 2, 2023, to February 1, 2026) for studies involving recombinant AAV.

## Data Availability Statement

The data and code required to generate the figures are available from https://github.com/deeprajs/AAV_impurity_separation. Additional data may be provided on request.

## Acknowledgments

S.M.H. acknowledges support from the William B. “Bill” Sturgis, ‘57 and Martha Elizabeth “Martha Beth” Blackmon Sturgis Distinguished Professorship in Chemical and Biomolecular Engineering.

