## Supporting Information for "IROP: Ionic Regulation of Permeation for High-Yield AAV Purification"

**Table S1.** Summary of AAV recovery and impurity removal in different experiments. AAV vector genome recovery from qPCR and total capsid recovery from ELISA include only the capsid concentrations in the final product stream following high salt filtration. Protein and dsDNA removal represents the fractions of these impurities present in the original feed that were not detected in the final product stream.

| Experiment | Salt (Conc.) | AAV vector genome recovery (qPCR) | AAV total capsid recovery (ELISA) | Protein (removal) | dsDNA removal |
|---|---|---|---|---|---|
| Static Salt Wash | NaCl (250 mM) | 0.25 | NM | NA | NA |
| Static Salt Wash | NaCl (250 mM) | 0.36 | 0.36 | Hemoglobin (NM) | NA |
| IROP | $MgCl_2$ (100 mM) | 0.76 | 0.95 | NA | NA |
| IROP | $MgCl_2$ (100 mM) | 0.74 | 0.84 | Hemoglobin (0.97) | NA |
| IROP | $MgCl_2$ (150 mM) | 0.93 | 1.16 | Hemoglobin (0.98) | NA |
| IROP | $MgCl_2$ (150 mM) | 0.96 | 0.89 | HCP (0.93) | 0.95 |

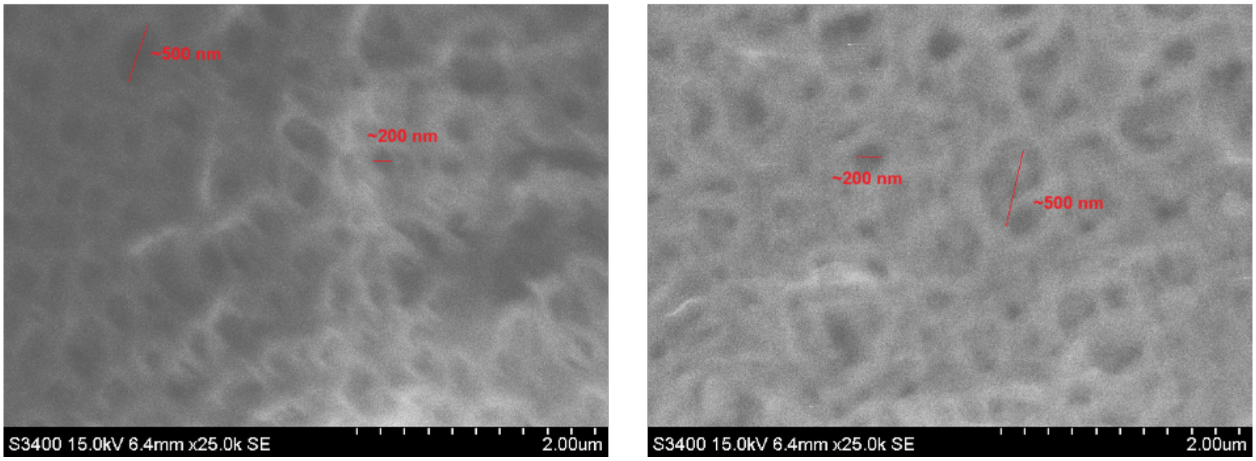


**Figure S1.** SEM images of (left) top and (right) bottom side of a nominal 30 nm PES membrane showing pores ranging from approximately 200 to 500 nm.
